# KLinterSel: Assessing Spatial Concordance among Selective Sweep Detection Methods, with an Application to the common cockle (*Cerastoderma edule*)

**DOI:** 10.1101/2025.08.21.671449

**Authors:** Antonio Carvajal-Rodríguez, Sara Rocha, Marina Pampín, Paulino Martínez, Armando Caballero

## Abstract

Detecting signals of positive selection, including selective sweeps, often involves applying several methods in parallel. Candidate regions identified by multiple approaches are commonly prioritized for further investigation, although agreement does not necessarily provide independent evidence of selection. Spatial concordance may arise by chance or reflect statistical dependencies among methods, shared demographic signals, or features of the genomic data such as linkage disequilibrium and heterogeneous marker density. To address this issue, we introduce two complementary statistical tests to evaluate whether spatial concordance among candidate sites exceeds expectations under their respective null models. The first is a fast parametric test based on a sequentially conditioned hypergeometric framework that evaluates k-way intersections among candidate sets at exact SNP positions or across genomic windows. The second relies on Monte Carlo resampling and compares observed inter-method distance profiles with those expected under random association, accounting for the empirical distribution of SNPs along the genome. By capturing different aspects of concordance, these approaches allow agreement among methods to be assessed across multiple spatial scales. Both tests are implemented in KLinterSel, which also identifies groups of candidate sites detected by different selection-detection methods within a user-defined distance threshold. We illustrate its application using simulated data, human chromosome 2 data from the 1000 Genomes Project, and candidate loci associated with resistance of the common cockle (*Cerastoderma edule*) to the parasite *Marteilia cochillia*. The software is written in Python and is available on GitHub, together with documentation and standalone executables for major operating systems.

## Introduction

At the genomic level, positive selection can alter the frequencies of neutral variants linked to beneficial alleles through genetic hitchhiking, leading to selective sweeps and reduced genetic diversity around selected sites (Caballero, 2020, Chap. 11; Maynard Smith & Haigh, 1974; Stephan, 2019). Numerous methods have been developed to detect selective sweeps (Panigrahi et al., 2023; Soni et al., 2023), and researchers may combine different approaches to capture complementary signatures of selection or account for differences in their sensitivity to evolutionary processes and confounding factors. Consequently, candidate sets need not overlap substantially (Schlamp et al., 2016), and a lack of agreement does not necessarily indicate poor performance of individual methods. Conversely, agreement does not necessarily provide independent validation, as concordant results may reflect methodological similarities or shared features of the underlying genomic data (Lotterhos, 2019; Lotterhos et al., 2018). Nevertheless, assessing concordance can still be valuable for prioritizing genomic regions for further investigation, particularly when methods targeting complementary signatures of selection identify nearby candidates or when genome-wide analyses yield large candidate sets.

Here, we introduce KLinterSel, a software tool implementing two complementary statistical tests to evaluate whether candidate SNPs identified by different methods show greater positional concordance along chromosomes than expected by chance. The hypergeometric k-intersection (HGkI) test uses a sequentially conditioned hypergeometric framework to assess k-way candidate overlap at exact positions or across genomic windows. The Kullback– Leibler-like Monte Carlo test (T_KL_) instead evaluates inter-method distance profiles by comparing observed distances among candidate SNPs with those expected from repeated permutations of SNP positions. Both tests provide *p*-values for assessing spatial concordance under their respective null models. KLinterSel also identifies candidate intersections within a user-defined genomic distance *D*. These analyses complement existing selection-detection methods but do not themselves detect selective sweeps or establish the biological validity of candidate loci.

We illustrate KLinterSel using high-density forward-time evolutionary simulations under different demographic and selection scenarios, human chromosome 2 data from the 1000 Genomes Project (YRI–CEU and GBR–CEU comparisons), and genomic and transcriptomic datasets from a previous study of resistance to the parasite *Marteilia cochillia* in the common cockle, *Cerastoderma edule*. Finally, we evaluate the false-positive rates and statistical power of HGkI and T_KL_ through controlled resampling experiments based on the genomic characteristics of the *Cerastoderma edule* datasets.

## Materials and Methods

### Full set of genomic positions

KLinterSel uses a reference set of genomic positions at which candidate loci can potentially be identified by the methods being integrated. Because all candidate sets analyzed here are SNP-based, we used the observed SNP positions as this reference set.

The two KLinterSel tests use these positions differently. HGkI does not involve positional resampling: for exact coincidences, the SNP positions define the universe of eligible units, whereas for regional overlap, only non-overlapping genomic windows containing at least one SNP are considered. The test evaluates overlap under a hypergeometric null model conditional on the number of eligible units and candidates identified by each method. By contrast, T_KL_ uses the observed SNP positions as its default resampling universe, thereby preserving the empirical spatial distribution of markers, including heterogeneous SNP density and regions with sparse coverage. Alternatively, the *--uniform* option allows T_KL_ to sample positions uniformly between the minimum and maximum observed SNP positions, without restricting them to actual SNP locations. More generally, KLinterSel is not intrinsically restricted to SNP-based candidates: other predefined sets of eligible genomic positions can be used when appropriate for the methods being compared.

### Hypergeometric k-way intersection (HGkI) test

The HGkI test evaluates whether candidate loci identified by two or more methods overlap more frequently than expected by chance. It is based on the hypergeometric distribution, which describes the overlap between subsets sampled without replacement from a finite universe, and extends this framework to multiple methods through sequential conditioning (Feller, 1968; Larson, 1982). The test considers either exact SNP coincidences or shared non-overlapping genomic windows of user-defined size. In the latter case, only windows containing SNPs are included, and multiple candidates from the same method within a window are counted once.

For example, consider 20 genomic units (SNP positions or SNP-containing windows), with methods A, B, and C identifying 4, 7, and 10 candidates, respectively. The hypergeometric distribution can be understood through the classical urn analogy. Consider an urn containing 20 balls. Under the null model, method A randomly draws 4 balls without replacement, marks them as its candidates, and returns them to the urn. Method B then independently draws 7 balls from the complete set of 20, also without replacement, marks them, and returns them to the urn. Finally, method C independently draws 10 balls from the same complete set. Thus, each method samples without replacement, but all 20 balls are available to every method, allowing the same genomic unit to be selected by multiple methods.

Suppose A and B share three units, two of which are also identified by C; HGkI evaluates whether this three-way intersection exceeds random expectations by considering all possible intermediate A-B overlaps, weighting each by its hypergeometric probability, and then evaluating the overlap with C conditionally. The same sequential procedure extends to any number of methods.

HGkI provides an exact right-tailed *p*-value under a null model of independent random assignment of candidates, conditional on the number of genomic units and candidates identified by each method. Let *K_k_* denote the size of the intersection among all *k* methods under the null model, and *K*_obs_ its observed value. The global *p*-value is then calculated as:

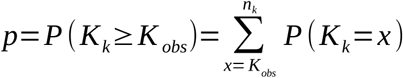

As a parametric test, it does not require Monte Carlo resampling. The complete mathematical derivation and computational details are provided in the Supplementary Material.

### Kullback-Leibler-like (T_KL_) Monte Carlo test

Unlike HGkI, which evaluates candidate overlap, T_KL_ assesses spatial concordance through the distribution of genomic distances between candidates identified by different methods. For each chromosome, all inter-method pairwise distances are calculated and sorted to obtain the observed distance profile. To establish random expectations, candidate positions are resampled without replacement from the available SNPs, preserving the number of candidates identified by each method. This procedure is repeated *R* times (10,000 by default), and the expected distance profile is obtained by averaging the corresponding ordered distances across replicates.

T_KL_ quantifies the discrepancy between the observed and expected distance profiles using a Kullback–Leibler-like statistic applied to vectors normalized to sum to one. Statistical significance is assessed by Monte Carlo resampling (Davison & Hinkley, 1997), considering only deviations toward shorter genomic distances: results are classified as non-significant when the observed median distance exceeds the expected median.

Because unequal candidate-set sizes may cause specific method pairs or individual methods to dominate the pooled distance profile, KLinterSel applies two safeguards when three or more methods have candidates. T_KL_ is not evaluated if a single method pair contributes more than 50% of all distances or if the distances involving one method exceed 95% of the total.

These restrictions do not apply to two-method comparisons. The complete mathematical formulation, resampling procedure, and computational details are provided in the Supplementary Material.

### Input data and software usage

#### Input files

KLinterSel requires at least three input files: one containing all analyzed SNP positions and two containing candidate SNPs identified by different selection-detection methods (one file per method). The first file may be provided in PLINK MAP (.map) format (Purcell et al., 2007), with four headerless columns (chromosome, SNP identifier, genetic position, and physical position), of which only the chromosome and physical position are used. Alternatively, CSV (.csv), TSV (.tsv), or plain-text (.txt) files with a header and two columns (chromosome and genomic position) can be used. Candidate files accept these same two-column formats or selscan .norm files. Additional methods can be included, subject only to available computational resources.

#### Intersection lists

The HGkI and T_KL_ tests identify chromosomes showing greater concordance among candidate sets than expected under their respective null models, based on overlap counts and inter-method distance profiles, respectively. Additionally, KLinterSel can identify groups of candidate positions detected by different methods within a user-defined physical distance threshold *D* (default *D* = 10 kb, adjustable using --*dist*). We refer to these groups as intersections. The choice of *D* should reflect the biological question, marker density, and expected extent of linked genomic signals. Users may also explore different thresholds to assess the sensitivity of the identified intersections.

When --*intersection* is specified, the program computes intersections in addition to the HGkI or T_KL_ test. To compute intersections only, without performing either test, use *–intersection --notest*. Because the distance criterion is applied between successive candidate positions, an intersection involving *k* methods may span up to (*k* − 1)*D*. Therefore, to restrict the maximum span of a *k*-way intersection to a desired distance *L*, the --dist parameter should be set to *L* / (*k* − 1).

#### Software usage

Standalone executables are provided for Windows, Linux, and macOS (arm64). Alternatively, KLinterSel can be installed from the source code available in the GitHub repository. By default, the hypergeometric test is executed; to run the Monte Carlo T_KL_ test, the --*TKL* flag can be specified. Detailed information on installation, usage, program options, and file formats, as well as an example dataset (test_dataset.zip), is provided in the program manual and on the project’s GitHub repository (https://github.com/noosdev0/KLinterSel/releases/latest).

### High-Density Forward-Time Evolutionary Simulations for the Evaluation of Selective Sweep Detection Methods and KLinterSel

High-density forward-time evolutionary simulations were performed in SLiM3 (Haller & Messer, 2019) to evaluate the XP-EHH and XP-nSL methods implemented in selscan v2.1 (Szpiech & Hernandez, 2014), the J_HAC_ method implemented in iHDSel v0.5.4 (Carvajal-Rodríguez, 2024), and the ability of KLinterSel to assess concordance among these methods.

Simulations modeled a 32-Mb chromosome with mutation and recombination rates of 10⁻⁸ per nucleotide per generation. An ancestral population of 20,000 diploid individuals was maintained for 9,700 generations before splitting into two populations of 10,000 individuals, followed by 300 additional generations. Six advantageous mutations (selection coefficient *s* = 0.1 in homozygotes, dominance coefficient *h* = 0.5) were introduced into one population at positions 1, 5, 10, 15, 20, and 30 Mb, with 20 initial copies of each mutation to reduce the risk of stochastic loss. Four scenarios were examined: constant population size, bottleneck followed by expansion, background selection, and their combination. The bottleneck involved a 20% population decline per generation for 10 generations, from 10,000 to 1,000 individuals for each population, followed by 25% growth per generation for another 10 generations, going back to 10,000 individuals. Background selection involved 5% deleterious mutations (selection coefficient *s* = -0.02 in homozygotes, dominance coefficient *h* = 0.5) throughout the simulation. Each scenario included a corresponding control without introduced advantageous mutations. Samples of 50 individuals per population yielded approximately 80,000 SNPs in the baseline scenario, with an average density of 2-3 SNPs/kb.

Independent Python-based resampling experiments assessed HGkI and T_KL_ false-positive rates and statistical power using parameters derived from the empirical *Cerastoderma* datasets. Null scenarios with different SNP spatial distributions and alternatives involving shared hotspots or reduced inter-method distances were considered. Full methodological details are provided in the Supplementary Material.

### Selection detection methods

The selection-detection methods considered in this study were chosen primarily to reflect the population-comparison framework that motivated the development of KLinterSel. For the simulated and human datasets, candidate loci were identified using three haplotype-based approaches specifically designed for comparisons between populations: XP-EHH (Sabeti et al., 2007) and XP-nSL (Szpiech et al., 2021) and J_HAC_. XP-EHH detects regions in which one population shows unusually long, high-frequency haplotypes relative to another population, whereas XP-nSL uses a related approach based on the number of segregating sites within haplotypes rather than genetic or physical distance. J_HAC_ provides a complementary signal by comparing the distribution of haplotype allelic classes (HAC), defined by their Hamming distance from a reference haplotype, between populations.

For the empirical *Cerastoderma edule* application, we additionally retained the candidate sets obtained with the population-differentiation approaches used in the original study of Pampín et al. (2023). For RAD-seq data, candidate SNPs were inferred using FDIST as implemented in ARLEQUIN v3.5 (Excoffier & Lischer, 2010), whereas for differentially expressed genes (DEGs) between different infection-level samples, F_ST_ was estimated using Genepop 4.7.5 (Rousset, 2008) and SNPs with the most extreme F_ST_ values were selected. These previously defined candidate sets are referred to here as Pampín23.

Thus, the methods considered provide complementary candidate sets based on haplotype structure and, for the *Cerastoderma* application, population differentiation. They were selected to address the population-comparison framework of the study rather than to constitute an exhaustive representation or benchmark of available methods for detecting selection. Details of the parameters used in each application are provided in the Data Processing section of the Supplementary Material.

### Normalization of selscan scores and window-based SNP collapse

XP-EHH and XP-nSL scores were standardized using either within-dataset normalization or reference-based normalization, with means and standard deviations estimated from the analyzed data or independent reference scores, respectively (Rahman et al., 2026; Szpiech, 2024). Reference scores were pooled across chromosomes for the empirical datasets or across 10 independent control replicates for each simulation scenario. Both approaches were evaluated in all three datasets. Within-dataset normalization was performed using the selscan companion program norm, whereas reference-based normalization was implemented using custom Python scripts.

To reduce the spatial redundancy of candidate SNPs in high-density datasets, we implemented a custom Python procedure for window-based SNP collapse, inspired by the selscan window-based outlier approach. Windows were ranked by enrichment in extreme scores, accounting for SNP density and considering both tails separately when applicable. The top 20% of eligible windows within each SNP-density class were selected, retaining only the most extreme SNP per selected window. This procedure was applied to the human and simulated datasets, but not to the lower-density *Cerastoderma* data. Full details are provided in the Data Processing Protocol section of the Supplementary Material.

### Empirical data

As empirical applications of KLinterSel, we analyzed human population-genomic data from the 1000 Genomes Project (The 1000 Genomes Project Consortium et al., 2015) and previously published population-genomic and transcriptomic data from the common cockle Cerastoderma edule associated with divergent selection in relation to the parasite Marteilia cochillia (Pampín et al., 2023) The selection-detection methods applied to each dataset are described above.

#### 1000 genomes project data

For the human empirical analysis, we used population-genomic data from the 1000 Genomes Project, focusing on chromosome 2 and, in particular, on the well-characterized MCM6/LCT region associated with lactase persistence (GRCh38/hg38: chromosome 2, position 135,851,076). Two population comparisons were considered: YRI versus CEU and GBR versus CEU. In addition to examining whether the selection-detection methods recovered a signal in the MCM6/LCT region, candidate overlap among methods was evaluated across the entire chromosome 2. The analyzed human data had a SNP density of approximately 2 SNPs/kb, comparable to that of the high-density simulated datasets.

#### Cerastoderma edule data

We will work with two types of data analysed by Pampín *et al*. *(2023)*: 6,077 anonymous SNPs from RAD-seq (restriction site-associated DNA sequencing) data and 13,004 SNPs located within differentially expressed genes (DEGs). The number of SNPs comes from filtering for invariant sites, missing values, and minimum allele frequency (MAF: see Supplementary Material for details on data filtering). The analyzed data had a SNP density of approximately one SNP every 120 kb for RAD-seq and one SNP every 60 kb for DEG data.

Samples exposed and unexposed to the marteiliosis outbreak were compared for the RAD-seq data, whereas different-level of infection individuals were compared for the DEG data (see Supplementary Material). Because XP-EHH, XP-nSL, and J_HAC_ are haplotype-based methods, both datasets were phased using SHAPEIT4 (Delaneau et al., 2019) with default parameters for low-density arrays.

For each data type (RAD-seq and DEG), four candidate sets were considered: the previously identified Pampín23 candidates and those obtained using XP-EHH, XP-nSL, and J_HAC_. The selscan methods were applied mostly with default parameters. For J_HAC_, fixed-size windows centered on outlier sites were applied without MAF filtering (MAF threshold = 0), consistent with the selscan analyses (see Data Processing Protocol in the Supplementary Material).

### KLinterSel analysis options

Unless otherwise stated, KLinterSel was run using its default redundancy-filtering criterion. Under this criterion, if two candidate sets contain comparable numbers of sites and at least 95% of the candidates in one set are located within 1 kb of a candidate in the other set, the redundant set is excluded from the analysis. The --*permissive* option disables this filtering step. For HGkI, rather than selecting a single window size, we evaluated candidate overlap across multiple spatial scales, from exact SNP-level coincidences (1 bp) to windows of 3 × 10³, 3 × 10⁴, 10⁵, 3 × 10⁵, 10⁶, 1.5 × 10⁶, and 2 × 10⁶ bp. This multiscale approach allows both exact coincidences and spatial concordance among nearby candidate sites to be assessed. For T_KL_, 10,000 permutations, corresponding to the default setting, were used to estimate the expected distance profile, except when the total number of inter-method distances exceeded 10 million, in which case 1,000 permutations were used.

Candidate lists containing more than 10,000 sites were analyzed using --*Kmax* 10000 together with a grouping distance of 1 kb (--dist 1000), thereby collapsing candidates from the same list that were separated by less than this distance. The --*paint* option was used to generate histograms of the observed and expected-by-chance distance profiles. The T_KL_ applicability safeguards described above remained in force even when redundancy filtering was disabled. When T_KL_ was not applicable because of pair or method dominance, KLinterSel reported the corresponding condition and recommended diagnostic analyses, such as evaluating the members of a dominant pair separately with the remaining methods or reducing an excessively large candidate set using --*Kmax* together with --*dist*. These follow-up analyses are diagnostic and do not replace a valid multimethod T_KL_ analysis.

Intersections were calculated separately by running KLinterSel with the --*intersection* and --*notest* options.

## Results

### High-Density Forward-Time Evolutionary Simulations

In the following analyses of simulated data, we focus on selscan results obtained using reference-based normalization followed by a window-based SNP-collapse procedure inspired by the window-based outlier approach implemented in selscan (Rahman et al., 2026; Szpiech, 2024). By identifying windows enriched in extreme normalized scores and retaining the top-scoring SNP from each selected window, this procedure substantially reduced spatial redundancy and facilitated the localization of selection signals. Window-based SNP collapse was applied only to the selscan results, as XP-EHH and XP-nSL produced large numbers of spatially clustered outlier SNPs, particularly under selective scenarios. J_HAC_ yielded substantially fewer candidate SNPs in both neutral and selective scenarios and was therefore analyzed without this additional filtering step. Figure 1 illustrates the ability of the different methods to recover the true selection targets under divergent selection, together with the signals detected in the neutral control. Detailed results before window-based SNP collapse, as well as comparisons with normalization based on the empirical distribution of the analyzed data, are provided in Supplementary Figure S1.

**Figure 1.**
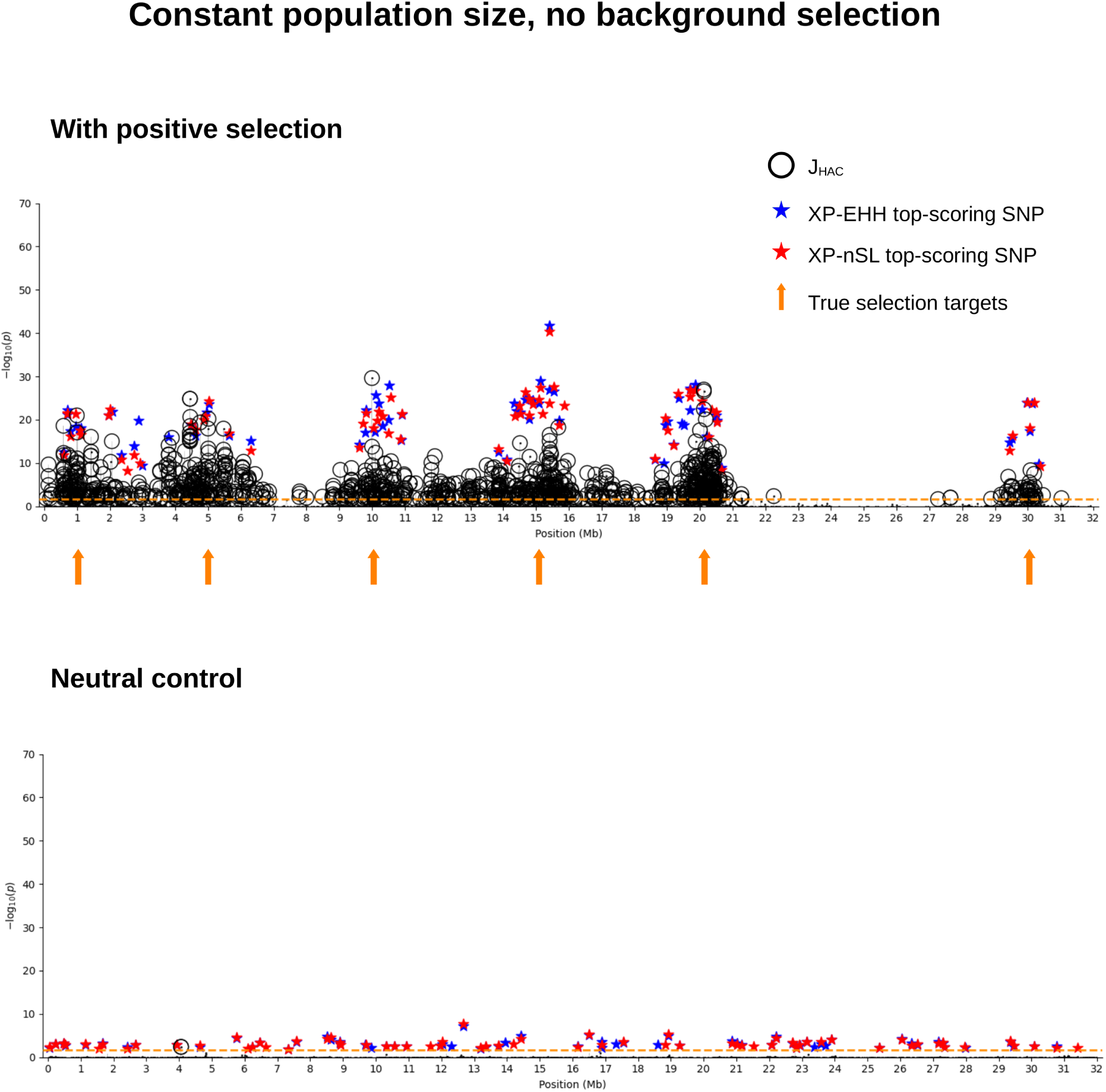
Selection signals along the simulated chromosome under the constant-population-size scenario without background selection, detected by XP-EHH (blue stars), XP-nSL (red stars), and J_HAC_ (black circles; fixed-size windows centered on outlier sites). The upper and lower panels show the scenario with positive selection and the corresponding neutral control, respectively. For the selscan methods (XP-EHH and XP-nSL), reference-based normalization followed by window-based SNP collapse was applied. The collapse procedure retains the highest-scoring SNP from each of the top 20% of eligible windows within each SNP-density class. Arrows indicate the six simulated selection targets at 1, 5, 10, 15, 20, and 30 Mb. The dashed horizontal line indicates the one-tailed selscan significance threshold (−log₁₀(0.0228), corresponding to *Z* > 2 and crit = 1 in selscan .norm files).

The neutral control provided an important contrast. For XP-EHH, before window-based SNP collapse, 1,994 of 84,868 SNPs exceeded the *Z* > 2 threshold, close to the 1,931 sites expected under the standard normal approximation. These outlier SNPs were distributed approximately uniformly along the chromosome. Following window-based SNP collapse, the number of retained candidate SNPs was reduced to 67. XP-nSL showed comparable results, whereas J_HAC_ identified only two significant candidate SNPs and did not produce a similar chromosome-wide background of outlier signals (Figure 1).

When additional selective scenarios involving background selection, a bottleneck followed by population expansion, or both were considered (Supplementary Figures S2–S4), stronger apparent selection signals were generally observed. Background selection also appeared to increase the number of additional signals surrounding the selected regions. The effects of normalization and window-based SNP collapse under these conditions are illustrated for the combined bottleneck-expansion and background selection scenario (Supplementary Figure S2), background selection alone (Supplementary Figure S3), and bottleneck-expansion alone (Supplementary Figure S4). Similar patterns of outlier detection in the absence of advantageous mutations were observed in the corresponding control simulations (Supplementary Figure S5).

Overall, both selscan methods identified numerous outlier SNPs even in the absence of advantageous mutations, whereas all three methods generally recovered the six selected regions under divergent selection. The marked increase in outlier SNPs under reference-based normalization in the selective scenarios highlights the usefulness of window-based SNP collapse for reducing spatial redundancy and providing a more concise representation of candidate selection signals in selscan results.

#### KLinterSel analysis: neutral and demographic controls

In the constant-size neutral scenario, HGkI was non-significant at all tested window sizes, and T_KL_ was not applicable to the three-method analysis because of the marked imbalance in candidate set sizes between the two selscan methods and J_HAC_. However, separate pairwise T_KL_ analyses of XP-EHH or XP-nSL with J_HAC_ yielded non-significant results. No three-way intersections were detected at either a 10-kb or a 0.1-Mb distance threshold.

In the control scenario with background selection, XP-EHH and XP-nSL each retained 69 candidate SNPs after window-based SNP collapse. HGkI detected significant three-way concordance only at window sizes of 0.1 and 0.3 Mb. The three-method T_KL_ analysis was prevented by the applicability safeguards, while separate pairwise comparisons with J_HAC_ were non-significant.

In the bottleneck-expansion control, XP-EHH and XP-nSL each retained 69 candidate SNPs after window-based SNP collapse, and HGkI was non-significant at all tested window sizes. T_KL_ was restricted to pairwise comparisons because of method dominance, and no significant concordance with J_HAC_ was detected. A similar pattern was observed in the combined bottleneck-expansion and background selection control.

Overall, KLinterSel detected no significant concordance in the constant-size neutral control. However, controls involving background selection occasionally yielded significant HGkI concordance. Detailed results obtained before window-based SNP collapse are provided in the Supplementary Material.

#### KLinterSel analysis: divergent selection scenario

In the constant-size selective scenario, J_HAC_ identified 1,155 candidate SNPs, whereas XP-EHH and XP-nSL each retained 68 candidates after window-based SNP collapse (Figure 1). HGkI detected significant concordance for exact SNP coincidences and across all tested window sizes. However, the three-method T_KL_ analysis and the pairwise comparisons involving J_HAC_ were not applicable because J_HAC_ contributed more than 95% of the pooled inter-method distances.

Regarding the genomic localization of candidate intersections, using a 10-kb distance threshold, three-way intersections among XP-EHH, XP-nSL, and J_HAC_ were found in the vicinity of all six simulated selection targets (1, 5, 10, 15, 20, and 30 Mb). For the most accurately localized targets, the closest intersections were approximately 30-50 kb from the true selected positions, whereas the largest deviations occurred for the selection targets at positions 15 and 20 Mb, with distances of approximately 0.2 and 0.5 Mb, respectively.

Increasing the intersection distance threshold to 0.1 Mb improved localization, with the closest intersections lying less than 1 kb from the true selected positions. However, additional intersections were also detected within approximately ±0.5 Mb of the selected targets. This contrasts with the constant-size neutral control, in which no three-way intersections were detected at either distance threshold.

The additional selective scenarios showed broadly similar patterns, although background selection appeared to increase both signal intensity and the number of additional signals surrounding selected regions (Supplementary Figures S2 and S3). Bottleneck-expansion generally produced stronger and more localized signals (Supplementary Figure S4). For example, in the combined bottleneck-expansion and background selection scenario (Supplementary Figure S2), XP-EHH and XP-nSL each retained 69 candidate SNPs after window-based SNP collapse, and neither candidate set was excluded by the redundancy filter. HGkI detected significant concordance at the SNP level and across all tested window sizes. Although the three-method T_KL_ analysis was prevented by J_HAC_ dominance, separate pairwise T_KL_ comparisons between J_HAC_ and each selscan method yielded significant results. In this scenario, three-way candidate intersections were confined to positions up to approximately 19 Mb at both the 10-kb and 0.1-Mb distance thresholds, as the window-based SNP-collapse procedure failed to retain the signal associated with the selected target at 30 Mb (Supplementary Figure S2, third panel).

### 1000 Genomes Project data: human chromosome 2

The analysis of human chromosome 2 using 1000 Genomes Project data showed that all three selection-detection methods recovered the well-known signal in the MCM6/LCT region associated with lactase persistence (GRCh38/hg38, chr2:135,851,076) in the YRI–CEU comparison. This signal was evident under reference-based normalization and remained detectable after window-based SNP collapse. In contrast, no comparable signal was observed in the GBR–CEU comparison (Figure 2). Beyond this specific region, we evaluated concordance among candidate sites identified by the three methods across the entire chromosome 2.

**Figure 2.**
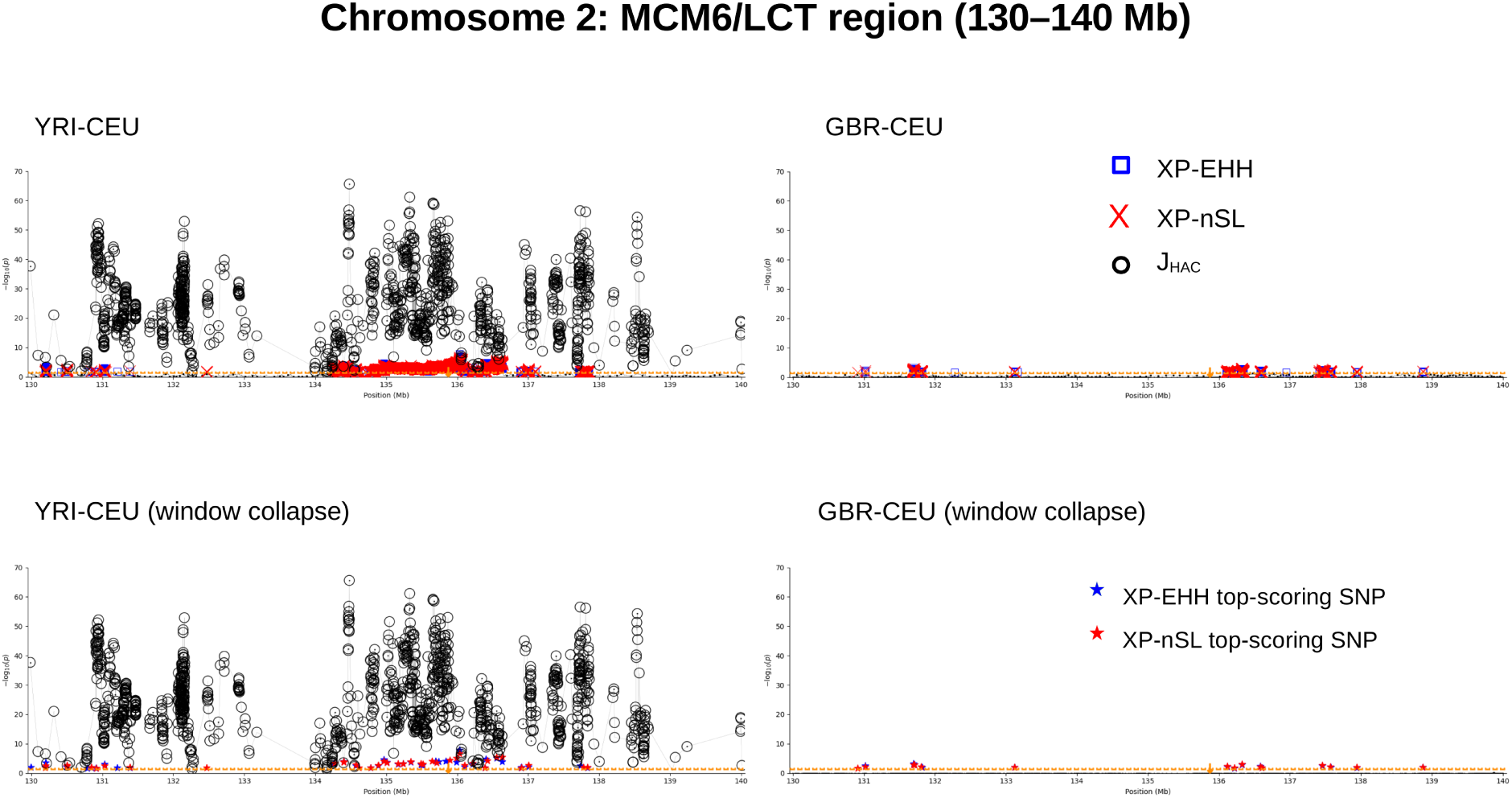
Selection signals in the MCM6/LCT region (130–140 Mb) of human chromosome 2, using 1000 Genomes Project data, detected by XP-EHH (blue squares), XP-nSL (red crosses), and J_HAC_ (black circles; fixed-size windows centered on outlier sites). Left and right panels show the YRI-CEU and GBR-CEU comparisons, respectively. Upper panels show reference-based normalization, whereas lower panels show the same normalization followed by window-based SNP collapse. Applied only to the selscan methods, this procedure retains one top-scoring SNP from each of the top 20% of eligible windows within each SNP-density class, indicated by blue (XP-EHH) and red (XP-nSL) stars. The dashed horizontal line indicates the one-tailed selscan significance threshold (-log10(0.0228)); *Z* > 2; crit = 1 in selscan norm).

#### YRI vs CEU

The two populations shared 610,081 SNPs, of which 22,696 were identified as candidates by XP-nSL, 22,952 by XP-EHH, and 13,407 by J_HAC_. Despite the large candidate sets, HGkI was applied directly and detected significant concordance for exact SNP matches and across all tested window sizes. For the T_KL_ analysis, candidates separated by less than 1 kb were first consolidated within each method, reducing the sets to 6,180, 4,957, and 10,080 candidates for XP-nSL, XP-EHH, and J_HAC_, respectively. Even with --*Kmax* filtering, the number of inter-method distances remained approximately 140 million. We therefore proceeded to window-based SNP collapse before applying T_KL_.

Window-based SNP collapse reduced the XP-nSL and XP-EHH candidate sets to 393 and 381 sites, respectively. This procedure was not applied to J_HAC_, which therefore retained its 10,080 candidates. The three-method T_KL_ analysis was prevented by the applicability safeguards because J_HAC_ dominated more than 95% of the inter-method distances. However, separate pairwise comparisons of J_HAC_ with each selscan method were significant.

Finally, analysis of the 1-kb-consolidated candidate sets identified 216 genomic positions involved in intersections within a 1-kb distance threshold. These included an extended cluster spanning approximately 135-136 Mb, encompassing the MCM6/LCT region and two SNPs flanking the lactase-persistence-associated variant rs4988235 (chr2:135,851,076), located 0.6 Mb and 0.18 Mb away, respectively.

#### GBR vs CEU

The two populations shared 837,888 SNPs, yielding 23,695 XP-nSL candidates, 24,182 XP-EHH candidates, and only 69 J_HAC_ candidates. HGkI detected significant concordance exclusively with 0.1- and 0.3-Mb windows, but not for exact SNP matches or smaller windows.

After consolidating candidates within 1 kb, the XP-nSL and XP-EHH sets were reduced to 4,183 and 4,047 sites, respectively, while J_HAC_ retained its 69 candidates. The three-method T_KL_ analysis was prevented by the pair-dominance safeguard, although separate pairwise comparisons of J_HAC_ with each selscan method were significant.

Subsequent window-based SNP collapse further reduced the XP-nSL and XP-EHH candidate sets to 507 and 496 sites, respectively. This procedure was not applied to J_HAC_, which therefore retained its 69 candidates. HGkI retained the same significance pattern, with significant concordance only for the 0.1-and 0.3-Mb windows. The three-method T_KL_ analysis remained inapplicable because of pair dominance, whereas both pairwise comparisons involving J_HAC_ were again significant.

Notably, inspection of the 1-kb-consolidated candidate sets (507 XP-nSL, 496 XP-EHH, and 69 J_HAC_ sites) revealed no three-way intersections anywhere along chromosome 2, even when the maximum distance between candidate SNPs was increased from 1 to 10 kb.

Overall, the YRI–CEU comparison showed stronger and more localized concordance among selection-detection methods, including the well-characterized MCM6/LCT region, whereas concordance in GBR–CEU was restricted to broader genomic scales in the HGkI analysis, with no closely spaced three-way candidate intersections detected.

### Genomic and transcriptomic data from the common cockle (*Cerastoderma edule*)

The analysis of all 19 chromosomes of *Cerastoderma edule* revealed broadly similar genomic patterns of selection signals in both the RAD-seq and DEG datasets under within-dataset and reference-based normalization, as illustrated for chromosome 18 in Figure 3. Below, we present the KLinterSel results obtained using reference-based normalization. Window-based SNP collapse was not applied because of the relatively low SNP density of these datasets.

**Figure 3.**
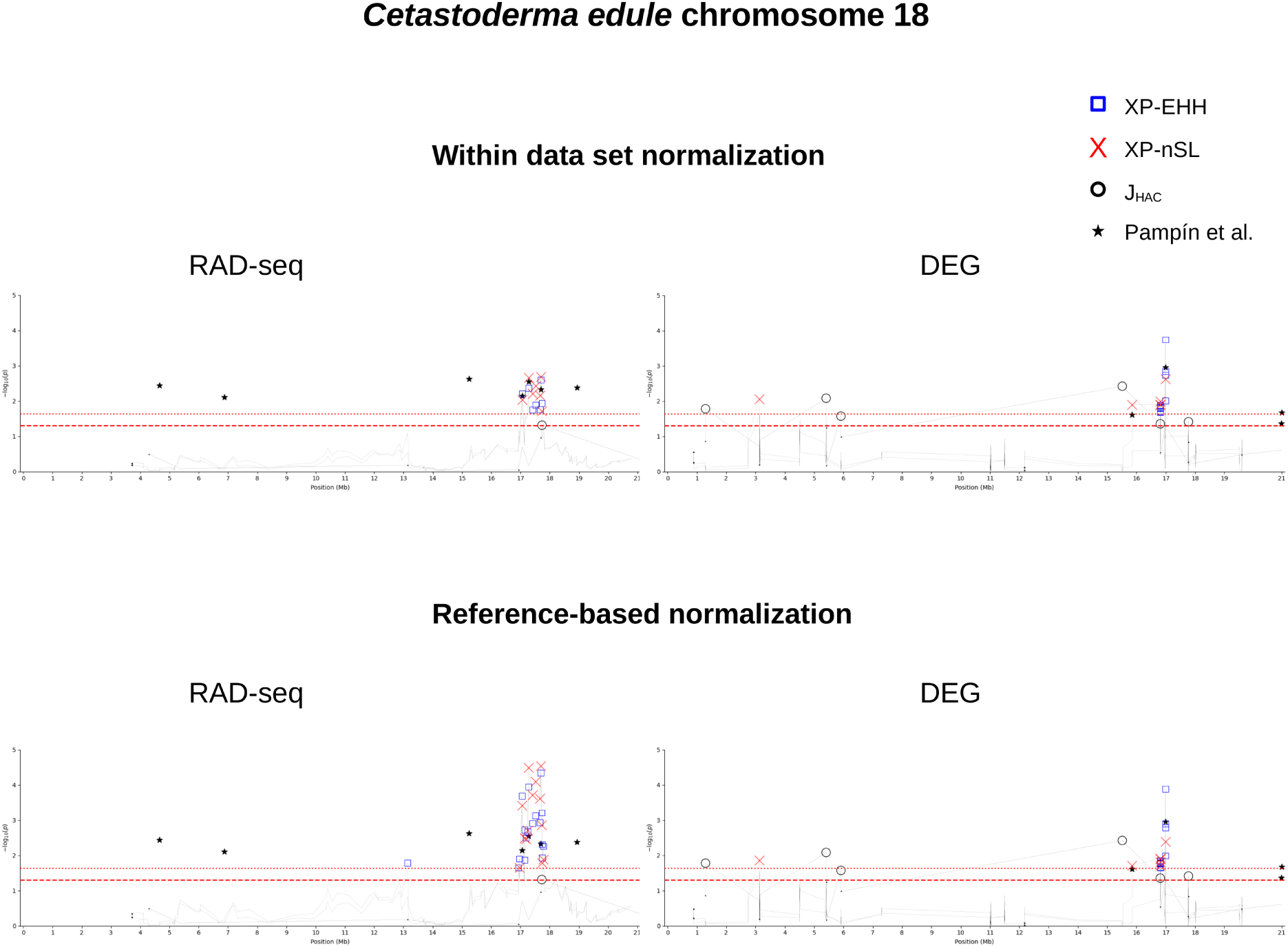
Selection signals along chromosome 18 of *Cerastoderma edule*, detected by XP-EHH (blue squares), XP-nSL (red crosses), and J_HAC_ (black circles; fixed-size windows centered on outlier sites). Black stars indicate significant SNPs previously reported by Pampín et al. (2023). Left and right panels show the RAD-seq and DEG datasets, respectively. Upper and lower panels show within-dataset and reference-based normalization of selscan statistics, respectively. The dotted horizontal line indicates the one-tailed selscan significance threshold (−log₁₀(0.0228); *Z* > 2; crit = 1 in selscan norm), and the long-dashed line indicates the J_HAC_ significance threshold (−log₁₀(0.05)).

#### RAD-seq data

For the RAD-seq dataset, a total of 6,077 SNPs distributed across 19 chromosomes were analyzed. Pampín et al. (2023) identified 153 candidate sites, whereas XP-EHH and XP-nSL detected 111 and 107 candidates, respectively, and J_HAC_ identified 54 candidates. The relatively limited number of candidates detected by J_HAC_, despite using the smallest possible fixed windows centered on outlier sites, is likely related to its reliance on linkage disequilibrium patterns. J_HAC_ requires a minimum level of linkage disequilibrium between adjacent sites to define informative windows, a condition that may be difficult to meet because of the sparse distribution of RAD-seq markers, with an average distance of approximately 120 kb between adjacent SNPs and linkage disequilibrium decaying to (*r*^2^ < 0.01) beyond 50 kb (Vera et al., 2022).

KLinterSel first showed that the degree of redundancy between the XP-EHH and XP-nSL candidate sets was below the exclusion threshold; therefore, neither set was removed from the analysis. The HGkI test detected no significant exact SNP overlap among the four candidate sets, but significant overlap was found on several chromosomes when window sizes of 100 kb or larger were used (Table 1). The T_KL_ test likewise detected significant spatial concordance among candidate sets on several chromosomes based on their inter-method distance profiles. Chromosomes 1, 6, 12, and 18 were significant under both HGkI and T_KL_ (Table 1). Chromosome 5 were found significant by HGkI but not by T_KL_. This discrepancy arises because the observed median distance exceeds that expected under the null model, leading T_KL_ to classify the chromosome as non-significant.

**Table 1.**
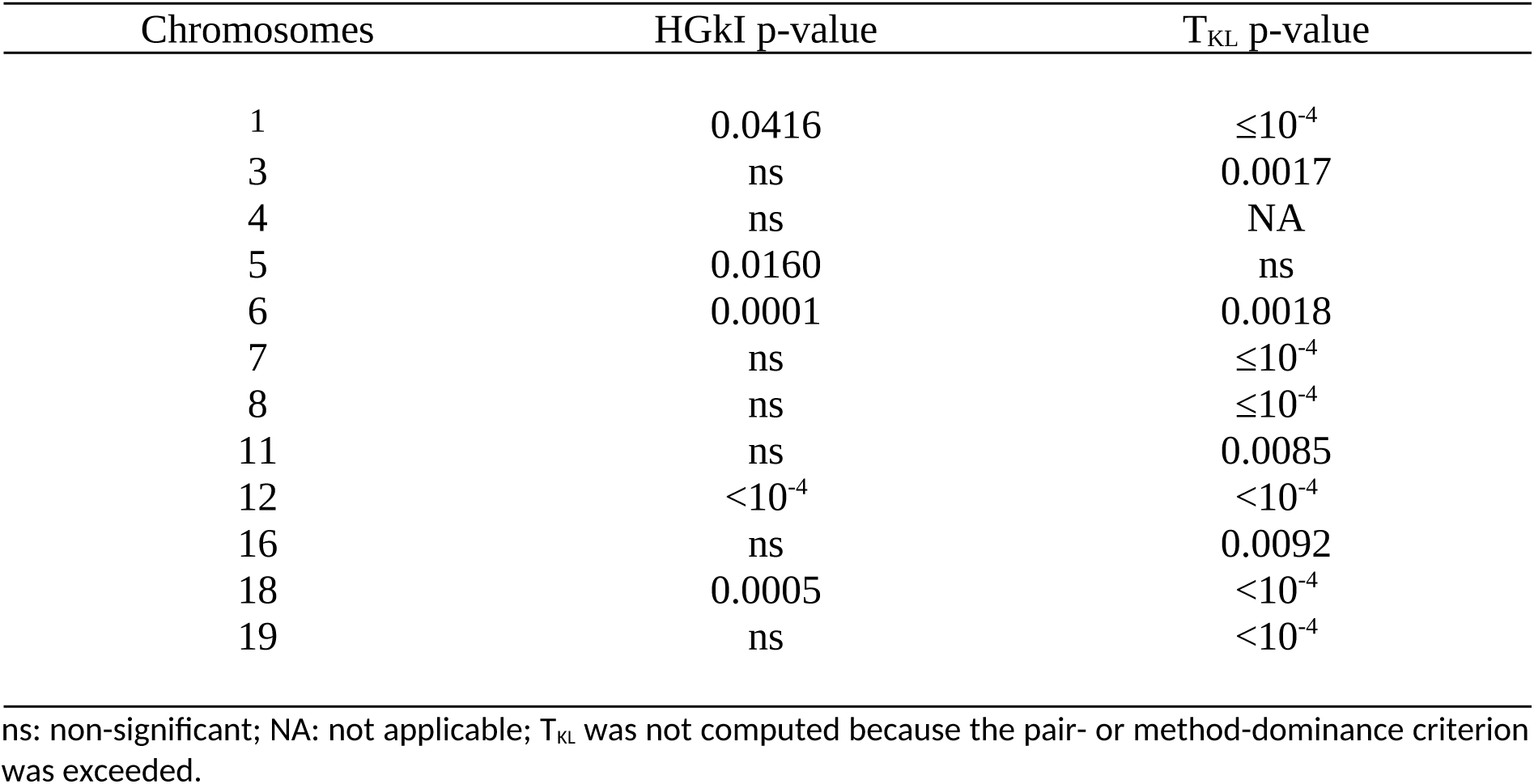
Chromosomes from RAD-seq data with significant overlaps between candidates of the four selection detection methods. The significance level is 0.05.

#### DEG data

For the DEG dataset, a total of 13,004 SNPs were analyzed. Pampín et al. (2023) identified 123 candidate sites, whereas XP-EHH, XP-nSL, and J_HAC_ identified 365, 326, and 191 candidates, respectively. KLinterSel showed that the redundancy between the XP-EHH and XP-nSL candidate sets was below the exclusion threshold; therefore, neither set was removed from the analysis. The HGkI test detected significant four-way concordance both at the exact SNP level and across all tested window sizes of 1 kb or larger. Chromosomes 6, 7, 10, 14 and 18 showed significant spatial concordance under both HGkI and T_KL_ (Table 2).

**Table 2.**
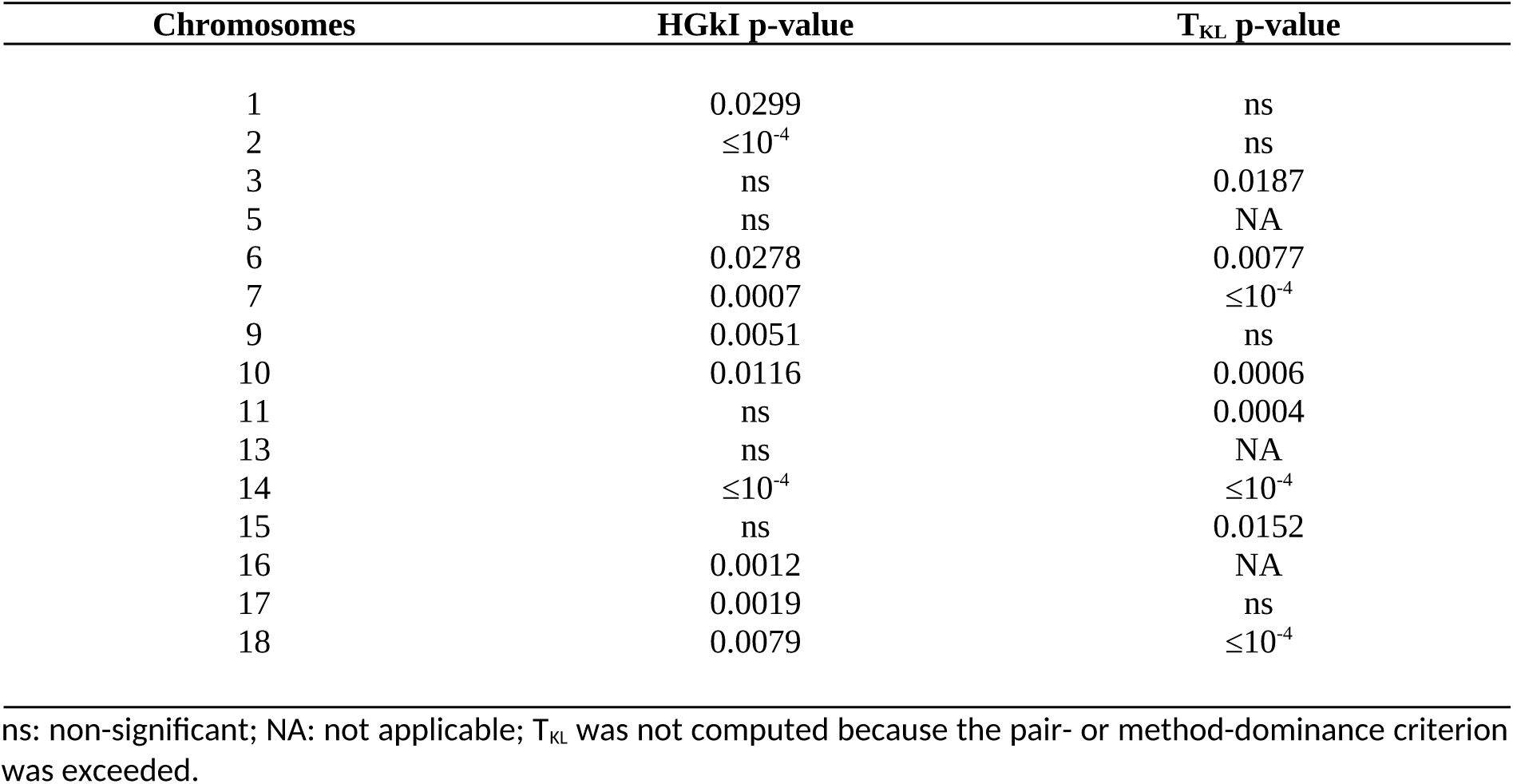
Chromosomes from DEGs data with significant overlaps between candidates of the four selection detection methods. The significance level is 0.05.

Chromosomes 1, 2 and 17 were detected by HGkI but not by T_KL_. As in the previous analysis, this discrepancy arose because the median of the observed inter-method distance profile was greater than that expected under the null model, causing T_KL_ to classify these chromosomes as non-significant.

#### Multiple testing correction

Null simulations preserving empirical genomic structure (Supplementary Material, Figure S6) showed that HGkI was conservative, with experiment-wide false-positive rates well below nominal expectations across spatial scenarios and window sizes. Further multiple-testing correction would therefore have unnecessarily reduced statistical power and was not applied. By contrast, T_KL_ showed good calibration, with an observed family-wise error rate close to theoretical expectations, supporting multiple-testing correction across chromosomes. Among the available procedures (Carvajal-Rodríguez, 2021) we used Myriads v1.4 (Carvajal-Rodríguez, 2018) to apply Holm’s sequential Bonferroni procedure (Holm, 1979), Benjamini-Hochberg FDR (Benjamini & Hochberg, 1995), the *q*-value approach based on positive false discovery rate estimation (Storey, 2003), and sequential goodness-of-fit procedure (SGoF, Carvajal-Rodríguez et al., 2009), all at a 5% threshold.

All chromosomes remained significant under the FDR and *q*-value criteria in both datasets. For RAD-seq, chromosome 11 became non-significant under Holm, and chromosome 16 under both Holm and SGoF. For DEG, chromosome 3 became non-significant under Holm and SGoF, and chromosomes 6 and 15 under Holm. All remaining chromosomes retained significance.

#### Candidate intersections between methods

KLinterSel computes candidate intersections across methods using *--intersection*, proceeding from pairwise to higher-order combinations. By default, intersections are calculated for chromosomes showing statistically significant concordance. The options *--intersection -- notest* allow analysis of all chromosomes, while adding *--chr-id* 18 restricts the analysis to chromosome 18. We focus on this chromosome because it was consistently supported by both HGkI and T_KL_ across the RAD-seq and DEG datasets, regardless of the normalization approach, and retained significance after multiple-testing correction. In contrast, chromosome 6 was supported by both tests in both datasets but became non-significant under Holm correction in the DEG dataset. Complete chromosome-level candidate counts, distance histograms, and HGkI and T_KL_ results are provided in the Supplementary Material.

Figure 3 reveals a region of candidate concordance around 17-18 Mb in both datasets. In RAD-seq, all four methods show a relatively precise overlap near 17.7 Mb. In DEG, one J_HAC_ candidate (SNP 17,772,723) falls within the same region, whereas the other methods identify signals shifted toward approximately 17 Mb. With a distance threshold of 0.1 Mb, KLinterSel identified six SNP positions (17,664,055; 17,697,950; 17,734,067; 17,740,229; 17,743,855; and 17,790,988) involved in four-method intersections in RAD-seq. In DEG, a threshold of 1 Mb was required to identify four-method intersections, involving SNPs spanning positions 15,514,725–17,772,723.

Figure 4 compares the observed (left panels) and expected (right panels) inter-method distance profiles for chromosome 18. The expected median distances slightly exceeded 6 Mb, compared with observed medians of 0.5 Mb (RAD-seq) and 1.1 Mb (DEG). For independently and uniformly distributed positions along a chromosome of length *L*, the theoretical median pairwise distance is *L*(1 − 1/√2) ≈ 0.293*L*, corresponding to approximately 5.9–6.5 Mb for chromosome 18 (*L* ≈ 20–22 Mb), consistent with the resampling results. Similarly, the theoretical proportion of distances ≤ 2 Mb is approximately 17-19%, compared with 10-15% in the expected profiles derived from genomic resampling and 30-35% in the observed profiles. These differences reflect the genomic structure preserved by resampling and the pronounced spatial clustering of candidate loci.

**Figure 4.**
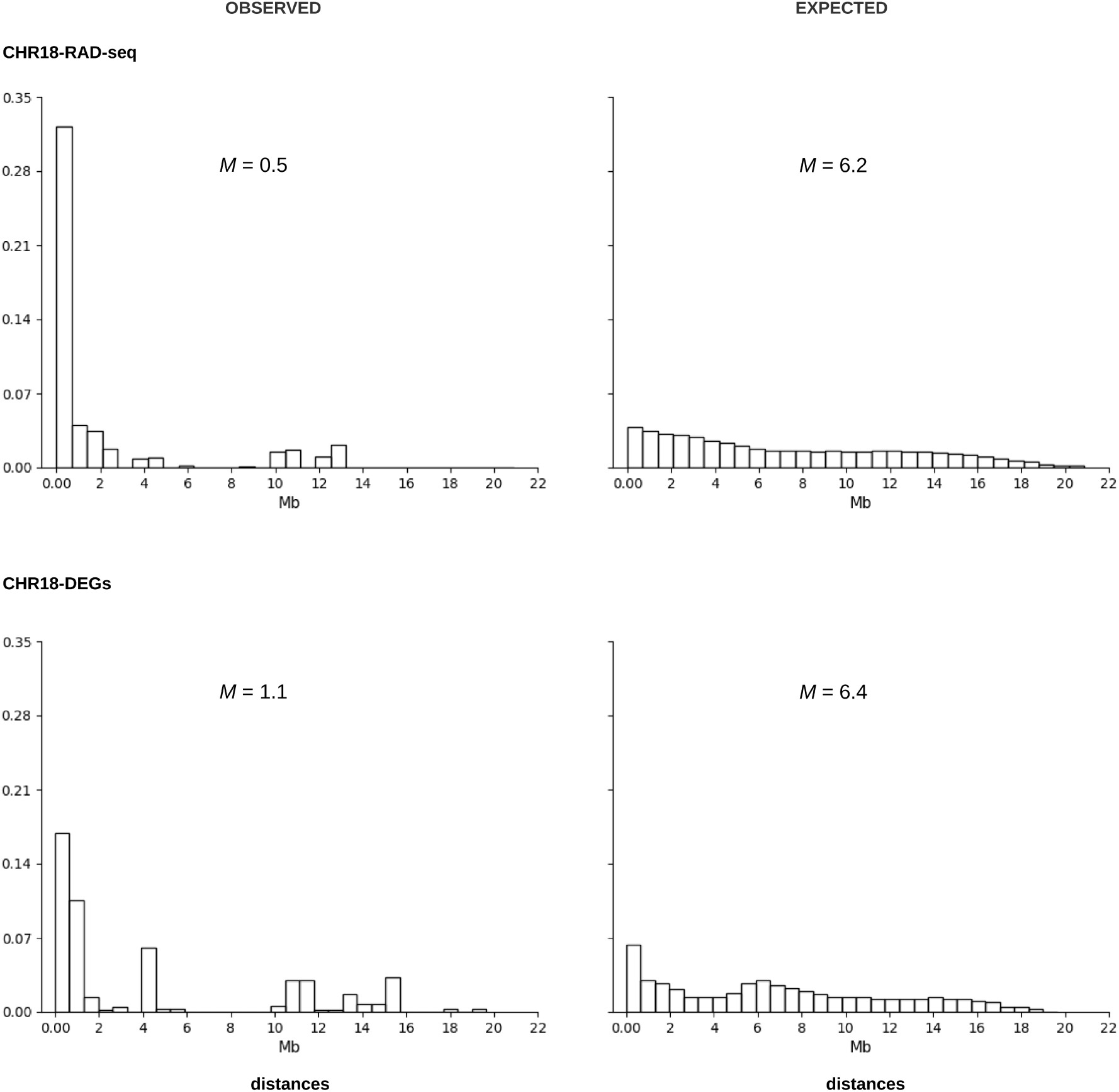
Histogram of observed (left) and expected by chance (right) distances between candidate sites for chromosome 18, obtained using four different selection detection methods. *M*: median measured in Mb units.

Additional power analyses showed that HGkI was particularly sensitive to localized candidate overlap, whereas T_KL_ performed comparatively better under more dispersed distance-compression scenarios (Supplementary Material, Figures S7–S8).

## Discussion

Although numerous methods are available for detecting selective sweeps (Panigrahi et al., 2023; Soni et al., 2023) similar genomic patterns can arise under different evolutionary scenarios (Johri et al., 2022; Soni et al., 2023; Soni & Jensen, 2024). Simulating plausible scenarios may help assess method reliability (Johri et al., 2022) but the underlying evolutionary histories are often unknown and difficult to model comprehensively. Consequently, multiple selection-detection methods are commonly applied to the same data (Horscroft et al., 2019), although they frequently identify different candidate loci (Schlamp et al., 2016). Even when methods agree, their concordance may reflect methodological similarities or shared features of the underlying genomic data rather than independent evidence of selection.

Assessing concordance becomes particularly challenging when large numbers of candidates are involved, especially because different methods may detect nearby but nonidentical SNPs within the same genomic region. KLinterSel addresses this problem through two complementary statistical tests that evaluate whether spatial concordance exceeds random expectations. HGkI assesses exact SNP coincidences or the co-occurrence of candidate SNPs from different methods within non-overlapping genomic windows. Applying the test across increasing window sizes allows concordance to be evaluated at different spatial scales. Larger windows may improve detection of nearby but nonidentical signals, although they also increase the probability of random overlap and eventually reduce spatial resolution.

Unlike HGkI, T_KL_ evaluates concordance through inter-method distance profiles. Expected profiles are generated by resampling candidate positions from the available genomic SNPs, thereby accounting for their nonuniform spatial distribution. This provides a more realistic null expectation than assuming uniformly distributed SNP positions, although KLinterSel also permits the latter assumption for comparisons with theoretical expectations. Thus, HGkI and T_KL_ provide complementary perspectives on candidate concordance, based on regional overlap and overall spatial proximity, respectively.

An important consideration is that agreement between closely related methods may reflect methodological redundancy rather than independent support for candidate loci. To limit this problem, KLinterSel excludes redundant candidate lists by default when they contain comparable numbers of candidates and at least 95% of candidates in one list lie within 1 kb of candidates in the other. T_KL_ also requires safeguards against disproportionate contributions to its pooled distance profile. When three or more methods have candidates, T_KL_ is not evaluated if a single method pair contributes more than 50% of all inter-method distances or if the distances involving one method account for more than 95% of the total. These restrictions help prevent misleading interpretations of multimethod concordance.

However, these safeguards do not address all potential sources of concordance. Although empirical-position resampling in T_KL_ preserves the spatial distribution of available SNPs, thereby indirectly accounting for some consequences of genomic heterogeneity, it does not explicitly model how variation in recombination rates, background selection, or demographic history affects the probability of identifying candidate loci. Consequently, shared responses of different selection-detection methods to these factors may still generate significant concordance in the absence of positive selective sweeps.

We illustrated the use of KLinterSel by assessing concordance among three selection-detection methods applied to high-density data from forward-time evolutionary simulations and human chromosome 2 data from the 1000 Genomes Project, and among four selection scans used to identify candidates for divergent selection in *Cerastoderma edule*.

### Concordance among Selective Sweep Detection Methods in High-Density Evolutionary Simulations

The usefulness of KLinterSel was illustrated using high-density genomic data simulated along a 32-Mb chromosome under four evolutionary scenarios: constant population size, constant population size with background selection, bottleneck followed by expansion, and bottleneck-expansion combined with background selection. Each scenario involved two populations, with positive selection acting at six genomic positions in one population, together with a corresponding control simulation without introduced advantageous mutations in either population.

Two main observations emerged from these simulations. First, in scenarios involving multiple selected sites, reference-based normalization of selscan statistics (Rahman et al., 2026) substantially increased the intensity and genomic extent of selection signals, making precise localization of the selected sites more difficult. This effect was particularly pronounced under bottleneck-expansion and/or background selection. Combining a window-based outlier approach, similar to that implemented in selscan (Rahman et al., 2026; Szpiech, 2024), with the retention of the highest-scoring SNP from each selected window substantially improved the localization of selective signals in subsequent KLinterSel analyses, although some weaker signals could be lost. Second, even in control simulations without introduced advantageous mutations, selscan identified large numbers of candidate SNPs after either within-dataset or reference-based normalization. Indeed, a substantial number of outliers is expected from the nominal significance threshold alone: under a standard normal distribution, the probability of Z > 2 is approximately 0.0228, corresponding to about 1,140 outliers among 50,000 SNPs. The distinction between individual extreme scores and spatially clustered selection signals has long been recognized and underlies window-based approaches to selection detection (Szpiech, 2024; Voight et al., 2006).

Because the absence of positive selection in a genomic region is generally unknown, such background signals complicate the identification of genuine selective sweeps. Although window-based SNP collapse greatly reduces the number of candidates, it does not eliminate them entirely. In this context, combining the reduced selscan candidate sets with J_HAC_ and evaluating their spatial concordance using KLinterSel helped eliminate most apparently spurious concordance signals in the control simulations, although some significant overlaps persisted, particularly under background selection.

Together, these findings suggest that KLinterSel, combined with appropriate candidate filtering and complementary selection-detection methods, can help distinguish concordant signals associated with positive selection from demographic and method-specific background signals in high-density genomic data.

### Concordance among selection scans using 1000 Genomes Project chromosome 2 data

The analysis of human chromosome 2 further illustrated the applicability of KLinterSel to large-scale genomic datasets. Although neither the YRI-CEU nor the GBR-CEU comparison represents a strictly neutral scenario, the well-established selective sweep associated with lactase persistence in the MCM6/LCT region provides a useful biological reference, with a strong signal expected in YRI-CEU but not in GBR-CEU. All three selection-detection methods successfully recovered this signal in the former comparison. However, genome-wide analysis involved more than 600,000 SNPs per comparison, with XP-EHH and XP-nSL each identifying over 20,000 candidate sites in both cases. In contrast, J_HAC_ identified approximately 13,400 candidates in YRI-CEU but only 69 in GBR-CEU, suggesting a markedly different response to these two population comparisons, potentially reflecting their contrasting levels of genetic differentiation.

Combining window-based SNP collapse for the selscan methods with KLinterSel analyses incorporating J_HAC_ provided further insight into the spatial concordance of selection signals. In YRI-CEU, HGkI detected significant concordance across all tested genomic scales, while intersection analyses identified signals in the MCM6/LCT region, approximately 0.18–0.6 Mb from rs4988235 after window-based SNP collapse, compared with less than 1 kb before collapse. This finding is consistent with previous evidence of intense recent positive selection associated with lactase persistence in European populations (Bersaglieri et al., 2004; Ingram et al., 2009). By contrast, HGkI concordance in GBR-CEU was restricted to broader genomic scales (0.1 and 0.3 Mb), with no three-way candidate intersections detected anywhere along chromosome 2.

These contrasting patterns illustrate the value of combining statistical tests of concordance with the genomic localization of candidate intersections when interpreting selection signals in large datasets.

### Concordance among selection scans on Cerastodema edule genomic data

Although XP-EHH and XP-nSL were expected to produce partially similar results, their overlap remained below the 95% redundancy threshold, and all four candidate sets were retained. We focused on chromosomes showing significant overlap and distance-profile patterns, as well as on genomic positions at which all methods coincided. Analyses were performed using both RAD-seq and DEG-associated SNPs (Pampín et al., 2023), with particular attention to chromosomes showing significant concordance in both datasets.

For several chromosomes, neither HGkI nor T_KL_ detected a significant departure from the corresponding null expectations. For other chromosomes, however, the concordance patterns were unlikely under the null model, indicating non-random agreement among methods. Chromosomes 6 and 18 were the only chromosomes detected by both HGkI and T_KL_ in both datasets regardless of the normalization approach. However, chromosome 6 became non-significant under Holm correction in the DEG dataset. In contrast, chromosome 18 retained significance after multiple-testing correction, with a strong concentration of selection signals around 17-18 Mb.

HGkI and T_KL_ provide complementary statistical evaluations of concordance. HGkI is a fast parametric test focused on regional overlap counts, whereas T_KL_ is a Monte Carlo approach based on full inter-method distance profiles. Consequently, T_KL_ may fail to detect chromosomes in which local clustering is offset by long-range dispersion, whereas HGkI may remain sensitive to localized overlap signals. Importantly, a lack of significant overlap or distance-profile deviation should not be interpreted as evidence for the absence of selection, but rather as a lack of evidence for non-random concordance among the methods examined. KLinterSel is therefore not intended to detect selection directly, but to provide a complementary framework for assessing whether agreement among candidate SNPs exceeds random expectations.

### Conclusions

KLinterSel provides a practical framework for assessing genomic concordance among selection-detection methods by combining HGkI tests of exact and regional overlap, T_KL_ analyses of inter-method distance profiles, and the identification of candidate intersections within user-defined distances. Applications to evolutionary simulations, human chromosome 2, and *Cerastoderma edule* demonstrate its usefulness for evaluating and prioritizing shared genomic signals in high-density datasets. In particular, simulations showed that the statistical tests and intersection analyses helped distinguish concordant signals around true selection targets from the numerous candidate signals detected under neutral control conditions. Although statistical concordance alone does not establish the evolutionary cause or biological validity of detected signals, KLinterSel provides a quantitative basis for prioritizing candidate regions for further investigation.

## Acknowledgements

We thank Enrique Santiago for providing the phased 1000 Genomes Project data. During the preparation of this work, the authors used ChatGPT (GPT-5.6) to review the style and clarity of the English. All AI-assisted text were subsequently reviewed and edited by the authors, who assume full responsibility for the content of the published article. This work was supported by the Marine Science Programme (ThinkInAzul) supported by the Ministerio de Ciencia e Innovación and Xunta de Galicia with funding from the European Union NextGenerationEU (PRTR-C17.I1) and European Maritime and Fisheries Fund, Xunta de Galicia (Grupo de Referencia Competitiva, ED431C 2024/22), Ministerio de Ciencia e Innovación (PID2022-137935NB-I00 and PID2025-168587NB-C21 – MICIU/AEI/10.13039/501100011033), Centro singular de investigación de Galicia accreditation 2024-2027 (ED431G 2023/07) and ERDF A way of making Europe.

## Data Accessibility and Benefit-Sharing section

### Data Accessibility Statement

The datasets generated during the current study, together with the SLiM simulation scripts and associated input files, are available in the Zenodo repository: [https://zenodo.org/records/16912703?preview=1&token=eyJhbGciOiJIUzUxMiJ9.eyJpZCI6IjM2ZTk2NWExLWE1MWMtNDg0My1iMmUwLTI2MmQzMDhkMDAxOCIsImRhdGEiOnt9LCJyYW5kb20iOiI5N2JmY2Q2ZGMwNzAyMjk5ODM4NTg2ZWQ1MjYyNmY1ZiJ9.RTsOg88slO8M82y2Ig_zmA5rH0r7LNmkWukiTTzktxa0Uk2BLCKN0IFBxwB9xirQZWoBXzMmI25bwwkNrNAjqA]

## Ethics statement

Not applicable. This study involved no new collection or experimentation on humans or animals and analysed only simulated and/or previously published data.

## Authors’ contributions

AC and ACR designed the study and conducted the simulations. MP, SR, PM and ACR prepared the data. SR, AC, and ACR analyzed the data. ACR developed the statistical tests and software and wrote the manuscript draft. All authors interpreted the data and results, and contributed to writing the manuscript. All authors read and approved the final manuscript.

